# The Retinal Basis of Light Aversion in Neonatal Mice

**DOI:** 10.1101/2022.01.20.477130

**Authors:** Franklin S. Caval-Holme, Marcos L. Aranda, Andy Quaen Chen, Alexandre Tiriac, Yizhen Zhang, Benjamin Smith, Lutz Birnbaumer, Tiffany M. Schmidt, Marla B. Feller

## Abstract

Aversive responses to bright light (photoaversion) require signaling from the eye to the brain. Melanopsin-expressing intrinsically photosensitive retinal ganglion cells (ipRGCs) encode absolute light intensity and are thought to provide the light signals for photoaversion. Consistent with this, neonatal mice exhibit photoaversion prior to the developmental onset of image vision, and melanopsin deletion abolishes photoaversion in neonates. It is not well understood how the population of ipRGCs, which constitutes multiple physiologically distinct types (denoted M1-M6 in mouse) encodes light stimuli to produce an aversive response. Here, we provide several lines of evidence that M1 ipRGCs that lack the Brn3b transcription factor drive photoaversion in neonatal mice. First, neonatal mice lacking TRPC6 and TRPC7 ion channels failed to turn away from bright light, while two photon Ca^2+^ imaging of their acutely isolated retinas revealed reduced photosensitivity in M1 ipRGCs, but not other ipRGC types. Second, mice in which all ipRGC types except for Brn3b-negative M1 ipRGCs are ablated, exhibited normal photoaversion. Third, pharmacological blockade or genetic knockout of gap junction channels expressed by ipRGCs, which reduces the light sensitivity of M2-M6 ipRGCs in the neonatal retina, had small effects on photoaversion only at the brightest light intensities. Finally, M1s were not strongly depolarized by spontaneous retinal waves, a robust source of activity in the developing retina that depolarizes all other ipRGC types. M1s therefore constitute a separate information channel between the neonatal retina and brain that could ensure behavioral responses to light but not spontaneous retinal waves.

## INTRODUCTION

Aversive responses to light in humans (McAdams et al., 2020; Noseda et al., 2010, 2019) and vertebrate animal models (Delwig et al., 2013; Johnson et al., 2010; Matynia et al., 2012; Zhang et al., 2017) are thought to originate from light-evoked signaling in melanopsin-expressing intrinsically photosensitive retinal ganglion cells (ipRGCs). While ipRGCs receive inputs from rods and cones (Güler et al., 2008), they are also photoreceptors themselves: melanopsin phototransduction is sufficient for photoaversion in adult *rd/rd cl* mice (Semo et al., 2010) and required for photoaversion in neonatal mice (Johnson et al., 2010), prior to the maturation of bipolar cell synapses with retinal ganglion cells on postnatal day 10 (P10) (Hoon et al., 2014).

What is the encoding of a light stimulus that evokes an aversive response? IpRGCs comprise multiple types, denoted M1-M6 in mice (Aranda and Schmidt, 2020; Do, 2019), with M1-M5 present in the neonatal retina (Caval-Holme et al., 2019; Lucas and Schmidt, 2019). M1-M6 differ in their light response properties, the molecules involved in their phototransduction, and the brain areas to which their axons project (Aranda and Schmidt, 2020; Do, 2019). Many of these brain regions are thought to relay light information for photoaversion (Okamoto et al., 2009; Routtenberg et al., 1978) and the negative effects of light on mood (An et al., 2020; Fernandez et al., 2018). Divergent axonal projections and previous demonstrations of type-specific contributions to non-image-forming functions (Chen et al., 2011; Rupp et al., 2019) raise the possibility that activity in a subset of ipRGC types could signal aversion.

The diversity of light responses across ipRGCs types is also apparent during development. Gap junction coupling of M2-M5 ipRGCs leads to functional classes that are distinct from anatomical types (Caval-Holme et al., 2019). Blockade of gap junction coupling or enhancement of coupling via reduced dopamine signaling leads to corresponding changes in the photosensitivity of M2-M5 ipRGCs (Arroyo et al., 2016; Caval-Holme et al., 2019; Kirkby and Feller, 2013). In contrast, M1 photosensitivity is invariant to modulation of gap junction coupling and therefore cell-autonomous (Caval-Holme et al., 2019). It is unknown how these developing circuits involving ipRGCs contribute to photoaversion.

Moreover, during the same developmental stage at which neonatal rodents exhibit photoaversion, the retina generates spontaneous activity patterns called retinal waves (Blankenship and Feller, 2010; Feller et al., 1996, 1997) that are thought to depolarize all RGCs (Ford et al., 2012), including ipRGCs (Kirkby and Feller, 2013). Excitation from retinal waves propagates along the axons of RGCs into the brain (Ackman et al., 2012; Mooney et al., 1996; Weliky and Katz, 1999) and seems poised to interfere with the detection of light that drives photoaversion. However, neonatal mice exhibit limited physical activity in the dark (Delwig et al., 2013; Johnson et al., 2010), indicating that retinal waves do not trigger a photoaversion-like behavior.

Here, we use a variety of approaches including transgenic mice and pharmacology to reveal which ipRGC types mediate photoaversion in neonatal mice. Additionally, we reveal that these ipRGCs exhibit reduced participation in retinal waves. Together, these data indicate that the signals underpinning photoaversion are reliable and informative.

## RESULTS

### Intensity threshold for photoaversion consistent with M1 or M3 photosensitivity

We first compared the light intensity threshold for photoaversion with the light intensity thresholds for the different types of ipRGCs, which vary over ~2 orders of magnitude (Caval-Holme et al., 2019; Lucas and Schmidt, 2019; Tu et al., 2005). To determine the light intensity threshold for photoaversion, we recorded movies of freely moving postnatal day 8 (P8) mice within a rectangular chamber, in darkness or with a beam of monochromatic blue light (λ_max_ = 470 nm; Full width at half-max = 29 nm; **Methods**) entering from one end (**Figure 1A**). We used automated tracking to measure head angle—as the maximum angle of deflection of the head away from the light source over time—and position within the chamber. The light was considered aversive if mice turned their heads and moved away from the light source (**Figure 1B**) (Delwig et al., 2013; Johnson et al., 2010; Jones et al., 2013; Mure et al., 2018). Photoaversion was absent in bilaterally enucleated mice (data not shown), consistent with a previous report in neonatal rats (Routtenberg et al., 1978), ruling out a role for melanopsin expression outside the eye (Delwig et al., 2018; Matynia et al., 2016; Sikka et al., 2014). Neonatal mice exhibited photoaversion at light intensities > 10^7^ photons μm^−2^ s^−1^ (**Figure 1D**), corresponding to ~10^5^ photons μm^−2^ s^−1^ at the retina, assuming a 100-fold loss of intensity through the closed eyelids (Tiriac et al., 2018). Ca^2+^ imaging and electrophysiology experiment in *ex vivo* retinas indicate that this light intensity reliably depolarizes neonatal M1s and M3s (Schmidt et al., 2008; Tu et al., 2005), but not M2s, M4s, and M5s (Caval-Holme et al., 2019).

**Figure 1:**
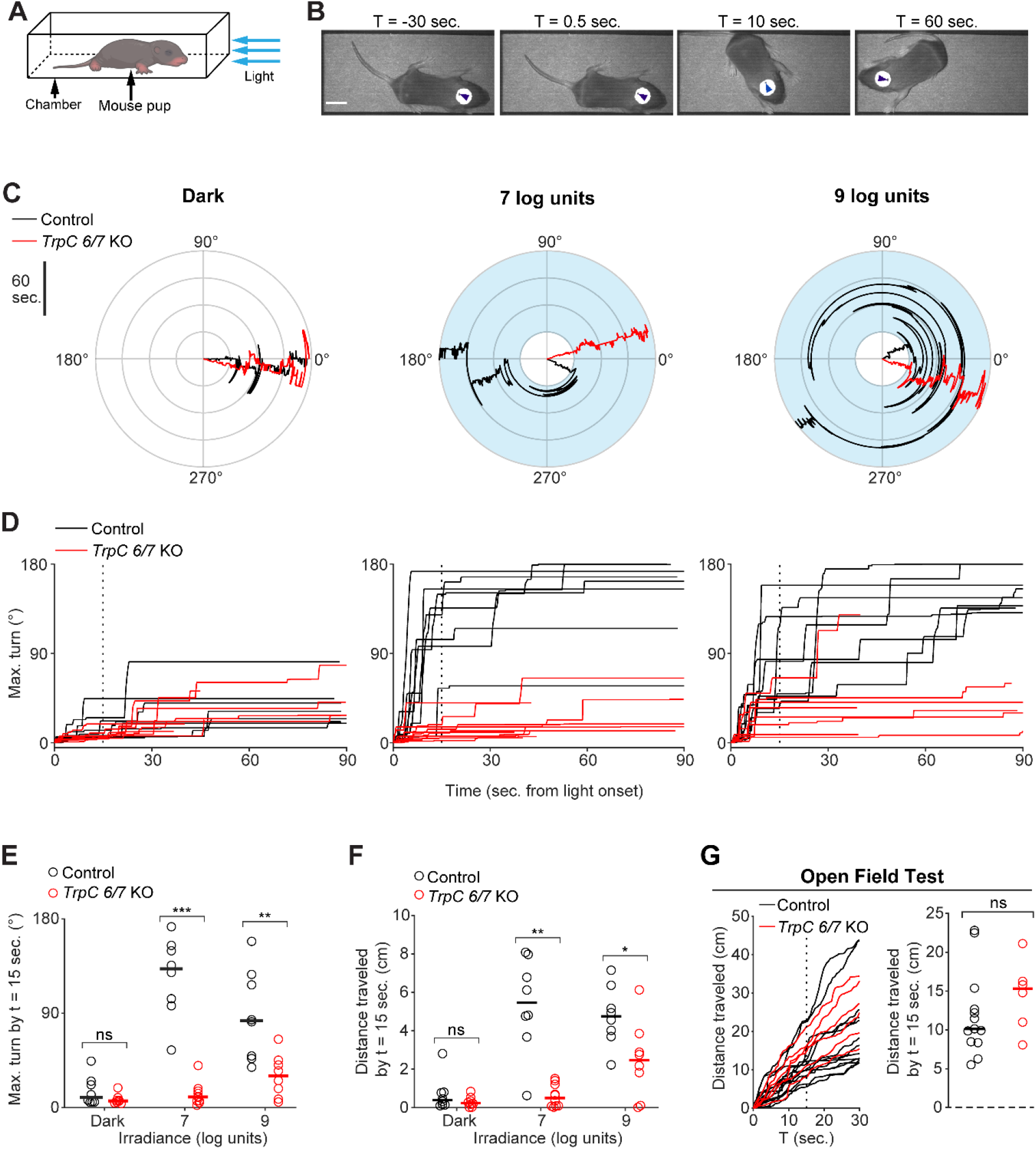
Photoaversion requires *TrpC 6/7*. A. Testing apparatus. Mouse pups (aged P 8, unless noted otherwise) were placed in a chamber. Trials began with a 30 sec. baseline in the dark. If the mouse remained oriented towards the LED (head angle < 45°), the trial proceeded for an additional 90 sec., either in darkness or with a light stimulus (blue arrows; λ_max_ = 470 nm; full width at half-maximum = 15 nm). Pups were monitored from above with infrared illumination (λ_max_ = 970 nm) to which melanopsin is ~10^12^ fold less sensitive (Emanuel et al., 2015). See Methods for further details. B. Video frames depicting the avoidance response of a P8 C57BL6/J mouse pup to a light stimulus (10^7^ photons μm^−2^ s^−1^; onset at t = 0 sec.). Arrowheads: automated tracking. Scalebar is 1 cm. C. Polar plots of head angle during representative behavioral tests of Control (black traces) or *TrpC 6/7* KO (red traces) mice in darkness (left) or with exposure to two different light intensities (middle and right; 7 log units = 10^7^ photons μm^−2^ s^−1^). Blue shading indicates the timing of the light stimulus. Time emanates outwards. D. Head turns of P8 genetic background control mice (Control, 1:1 C129:C57BL6/J; black traces) and mice lacking TRPC6 and TRPC7 ion channels (*TrpC 6/7* KO; red traces), as a function of light intensity. Vertical dashed line: time sample (15 sec.) used for statistical comparisons in E. and F. Each mouse was tested in darkness and at both light intensities. n = 8 mice from each group. E. Comparison of head turns of Control and *TrpC 6/7* KO mice, sampled 15 sec. after light onset. Horizontal bars are medians. ns: not significant (p = 0.19), **p < 0.05, ***p < 0.0005, Wilcoxon rank-sum test. Same mice as in D. F. Comparison of movement (travel distance of the centroid of the infrared-reflective sticker) in Control and *TrpC 6/7* KO mice, sampled 15 sec. after light onset. Horizontal bars are medians. ns: not significant (p = 0.51), *p < 0.05, **p < 0.005, Wilcoxon rank-sum test. n = 8 mice from each group. Note: some trials were excluded due to head-tracking errors. Same mice as in D. and E. G. An open field test indicated no difference between locomotion in Control (black traces) and *TrpC 6/7* KO (red traces) mouse pups tested at P11. Left: travel distance during the open field test. Vertical dashed line indicates the time sample (15 sec.) used for statistical comparison. Right: comparison of locomotion Horizontal bars are medians. ns = not significant. Wilcoxon rank-sum test, p = 0.28. N = 13 Control mice and 6 *TrpC 6/7* KO mice.

### Photoaversion requires *TrpC 6/7*

To identify the type of ipRGCs that mediate photoaversion we first sought to identify the phototransduction mechanisms involved in the behavior. To this end, we measured the behavior in mice that lack TRPC6 and TRPC7 ion channels (*TrpC 6/7* KO), the primary phototransduction channels in M1 ipRGCs (Jiang et al., 2018; Sonoda et al., 2018; Xue et al., 2011). Strikingly, P8 *TrpC 6/7* KO mice had no detectable response to the threshold intensity for WT mice (10^7^ photons μm^−2^ s^−1^; **Figure 1D, 1E**). In response to a saturating intensity (10^9^ photons μm^−2^ s^−1^), *TrpC 6/7* KO mice made small head turns and movements (**Figure 1E, 1F**; 2/8 *TrpC 6/7* KO mice turned more than 45° within 90 seconds of light onset, compared to 9/9 genetic background control mice. For videos of the behavior, see http://fellerlab.squarespace.com/movies). Dramatically reduced photoaversion in *TrpC 6/7* KO mice did not result from a general movement deficit, as *TrpC 6/7* KO and control mice traveled similar distances during an open field test (**Figure 1G; Methods**). TRPC6/7 ion channels are therefore critical for photoaversion, with remaining phototransduction mechanisms enabling residual behavioral responses to a saturating light intensity.

### M1 ipRGCs of *TrpC 6/7* KO mice exhibit reduced photosensitivity

In adult *TrpC 6/7* KO mice, M1s lack virtually all photocurrent (Xue et al., 2011), while some M2s lack a component of their photocurrent and M4s are unaffected (Jiang et al., 2018; Sonoda et al., 2018). However, the role of TRPC6/7 in developing ipRGCs is not known. To characterize the neonatal retina’s population light response in the absence of TRPC6/7 phototransduction, we used two-photon Ca^2+^ imaging to measure light-evoked responses from individual neurons within the ganglion cell layers of retinas from P7-9 WT and *TrpC 6/7* KO mice. We presented a range of light intensities (10^4^ −10^8^ photons μm^−2^ s^−1^; 420 or 470 nm; **Methods**) (**Figure 2A**) corresponding to ~10^6^ −10^10^ photons μm^−2^ *s*^−1^ *in vivo* during the photoaversion assay. To isolate light-evoked Ca^2+^ transients from those triggered by retinal waves, we superfused retinas with a solution containing the nicotinic acetylcholine antagonist Dihydro-β-erythroidine (DHβE; 8 μM (Bansal et al., 2000). The retinas of *TrpC 6/7* KO mice contained light-responsive cells, but these were present at a lower density (**Figure 2B**), consistent with a subset of ipRGCs dependent on TRPC6/7 for phototransduction.

**Figure 2:**
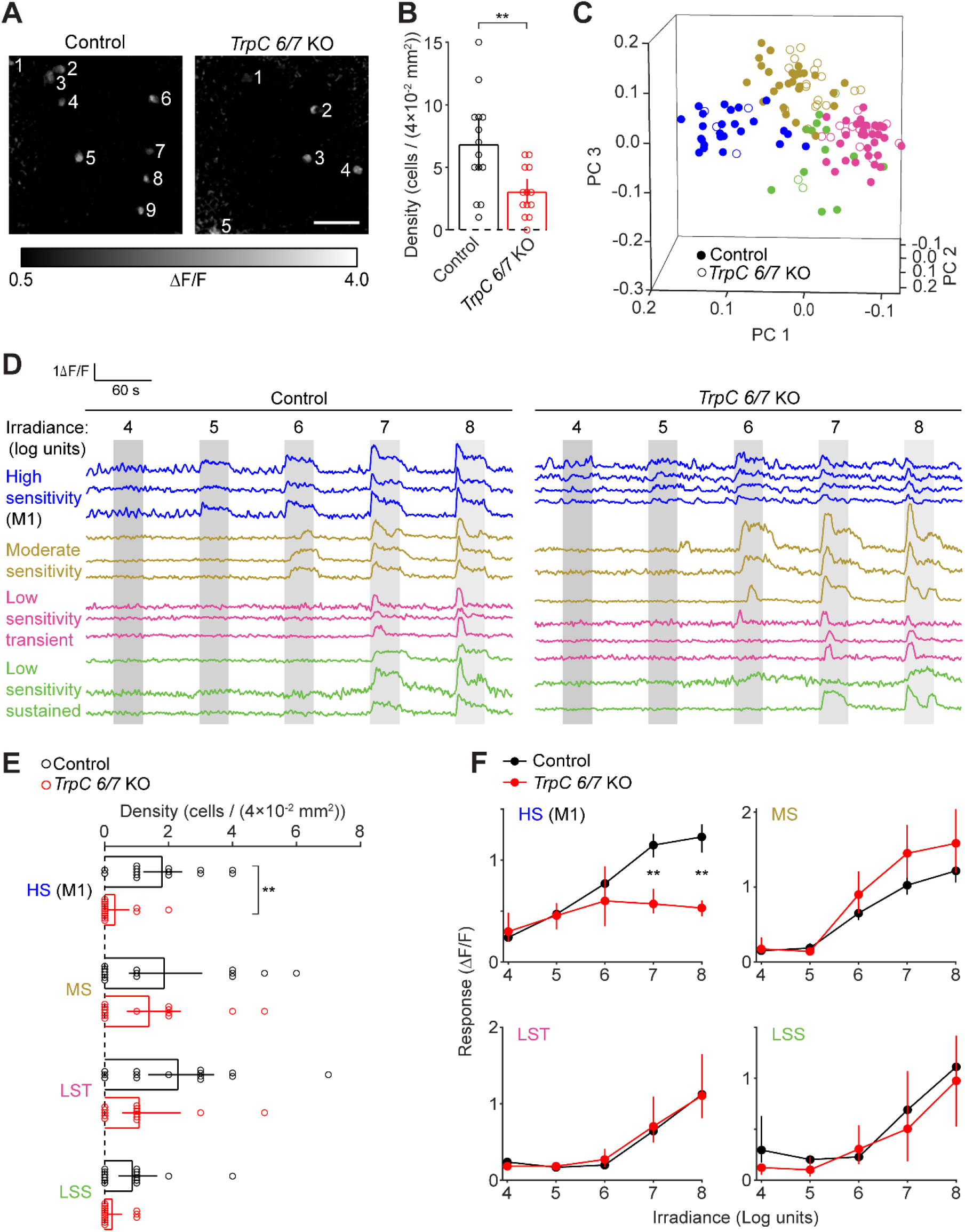
M1 ipRGCs of *TrpC 6/7* KO mice exhibit reduced photosensitivity. A. Fluorescence responses to the presentation of a light stimulus (λ_max_ = 470 nm, 10^8^ photons μm^−2^ s^−1^) observed with two-photon imaging of the Ca^2+^ indicator Cal 590 AM. Representative fields of view (FOV, 200 × 200 μm) are depicted for a retina from a wildtype control mouse (left panel from a matched genetic background (1:1 C129:C57BL6/J) and a *TrpC 6/7* KO mouse (right panel). Numbering indicates light-sensitive cells. Scale bar: 50 μm. B. Comparison of the number of light-sensitive cells per FOV. Error bars are bootstrapped 95% confidence intervals. **p < 0.005, two-sample t test. Control: n = 14 FOV from 5 mice. *TrpC 6/7* KO: n = 13 FOV from 4 mice. C. Visualization of each cell’s light-evoked fluorescence transient within the space of the first three principal components of population activity. Each point represents a cell from the retina of a Control (closed symbols) or *TrpC 6/7* KO mouse (open symbols). Point colors correspond to functional clusters named in D. D. Fluorescence traces (ΔF/F) from representative cells of each functional cluster recorded in the retinas of Control (left panel) and *TrpC 6/7* KO (right panel) mice. Functional clusters are named and color-coded according to the sensitivity (irradiance threshold) and duration of their light responses. Grey bars indicate the timing of light stimuli. 4-8 log units: 10^4^-10^8^ photons μm^−2^ s^−1^. E. Comparison of irradiance-response relations between control and *TrpC 6/7* KO cells within each functional cluster. **p < 0.005, Wilcoxon signed-rank test. Control: n = 95 cells (with 24, 30, 29, and 12 cells assigned to HS, MS, LST, and LSS, respectively). *TrpC 6/7* KO: n = 39 cells (with 4, 14, 19, and 2 cells assigned to the functional clusters listed above). F. Comparison of the number of cells within each functional cluster, per FOV, in Control and *TrpC 6/7* KO retinas. **p < 0.005. ns: p > 0.05. Wilcoxon signed-rank test. Same cells as in E.

To determine which ipRGC types have disrupted photosensitivity in *TrpC 6/7* KO retinas, we first computationally classified cells according to their light-evoked Ca^2+^ transients (**Figure 2C, 2D**) (Baden et al., 2016; Caval-Holme et al., 2019). We identified four functional types of cells in the WT retinas, defined by the threshold and duration of their light responses: High Sensitivity (HS), Moderate Sensitivity (MS), Low Sensitivity Transient (LST), and Low Sensitivity Sustained (LSS). We then used the model that identified functional types in the WT to classify light-responsive cells in the *TrpC 6/7* KO retinas (**Methods**). Notably, *TrpC 6/7* KO retinas contained cells of each functional type found in the WT though with a lower densities of HS cells (**Figure 2F**). Moreover, compared to HS cells in the WT, the HS cells in the *TrpC 6/7* KO had similar light-intensity thresholds but smaller amplitude Ca^2+^ transients (**Figure 2E**). Irradiance tuning of Ca^2+^ transients in other functional types (MS, LST, and LSS) was unaffected. Hence, we hypothesize that the reduced light sensitivity of the population of HS cells in the *TrpC 6/7* KO is insufficient to drive photoaversion at these lower light intensities.

The light-evoked Ca^2+^ transients of HS cells closely resembled those previously reported for neonatal M1s (Caval-Holme et al., 2019). To determine if HS cells corresponded to M1s, we anatomically identified M1s after Ca^2+^ imaging by imaging their dendrites (which stratify exclusively in the OFF sub-lamina of the inner plexiform layer), either in post-fixed retinas, using melanopsin immunohistochemistry (data not shown), or in the living *ex vivo* retinas of mice that express GFP under the control of the *Opn4* gene (Opn4::eGFP) (Caval-Holme et al., 2019; Schmidt et al., 2008). Our unsupervised classification closely reflected the anatomy: 6/7 of the anatomically identified M1s were assigned to the HS functional type, with the remaining M1 assigned to the MS functional type. Thus, reduced photoaversion in *TrpC 6/7* KO mice corresponded with a specific reduction in M1 photosensitivity.

### Normal photoaversion in mice with only M1 ipRGCs

To determine if signaling from M1 ipRGCs alone is sufficient for photoaversion, we measured the behavior in Opn4^Cre^ Brn3b^DTA^ mice (DTA), in which Cre-dependent expression of diphtheria toxin under the control of the Brn3b promoter ablates all ipRGCs by P7 except a subset of ~400 M1s that lack Brn3b expression (Chen et al., 2011; Chew et al., 2017). Remarkably, DTA mice exhibited photoaversion indistinguishable from littermate controls (**Figure 3**), despite their lack of M2-M6 ipRGCs. Thus, signals originating from a small subset of M1 ipRGCs are enough to drive photoaversion over a hundred-fold range of light intensity.

**Figure 3:**
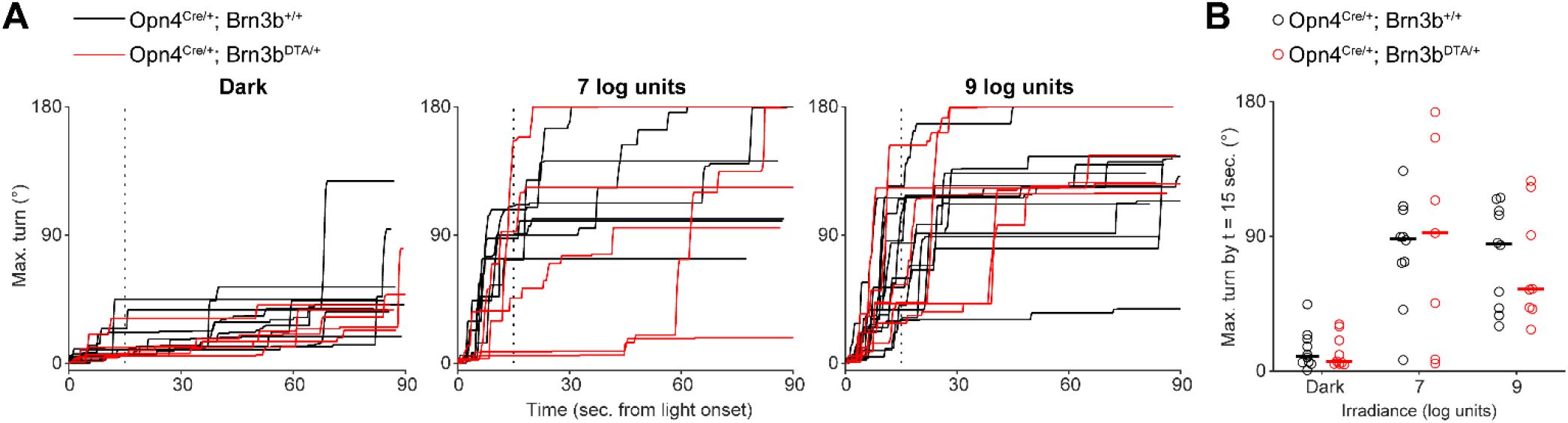
Normal photoaversion in mice with only M1 ipRGCs. A. Head turns in littermate Opn4^Cre^ mice (Control; black traces) and Opn4^Cre^ Brn3b^DTA^ mice with diphtheria toxin ablation of all ipRGCs save for ~400 M1s (DTA; red traces). 7 or 9 log units: 10^7^ or 10^9^ photons μm^−2^ s^−1^. Each mouse was tested in darkness and at both light intensities. B. Comparison of turns by Control and DTA mice, sampled 15 sec. after light onset (vertical dashed line in A). Horizontal bars indicate medians. There were no significant differences between turns by Control and DTA mice (p = 0.78, 0.74, and 0.97 for Dark, 7 log units, and 9 units, respectively. Wilcoxon rank sum test). n = 10 Control and 8 DTA mice. Note that some mice were excluded from (A) due to errors in automated tracking.

### Photoaversion does not require gap junction coupling in the retina

Our previous studies indicate that non-M1 ipRGCs are extensively gap junction coupled and that blockade of gap junction coupling significantly diminishes the photosensitivity of M2-M6, but not M1s (Caval-Holme et al., 2019) (**Figure 4A**). Hence, to investigate whether signaling from the non-M1 ipRGCs contributes to photoaversion, we assayed photoaversion in mice that received intraocular injections of the gap junction antagonist Meclofenamic Acid (MFA; ~70 μM in the vitreous). Mice injected with MFA exhibited photoaversion behavior indistinguishable from saline-injected littermate controls (**Figures 4B and C**), suggesting that disruption of photosensitivity in M2-M6, but not M1, had little effect on photoaversion.

**Figure 4:**
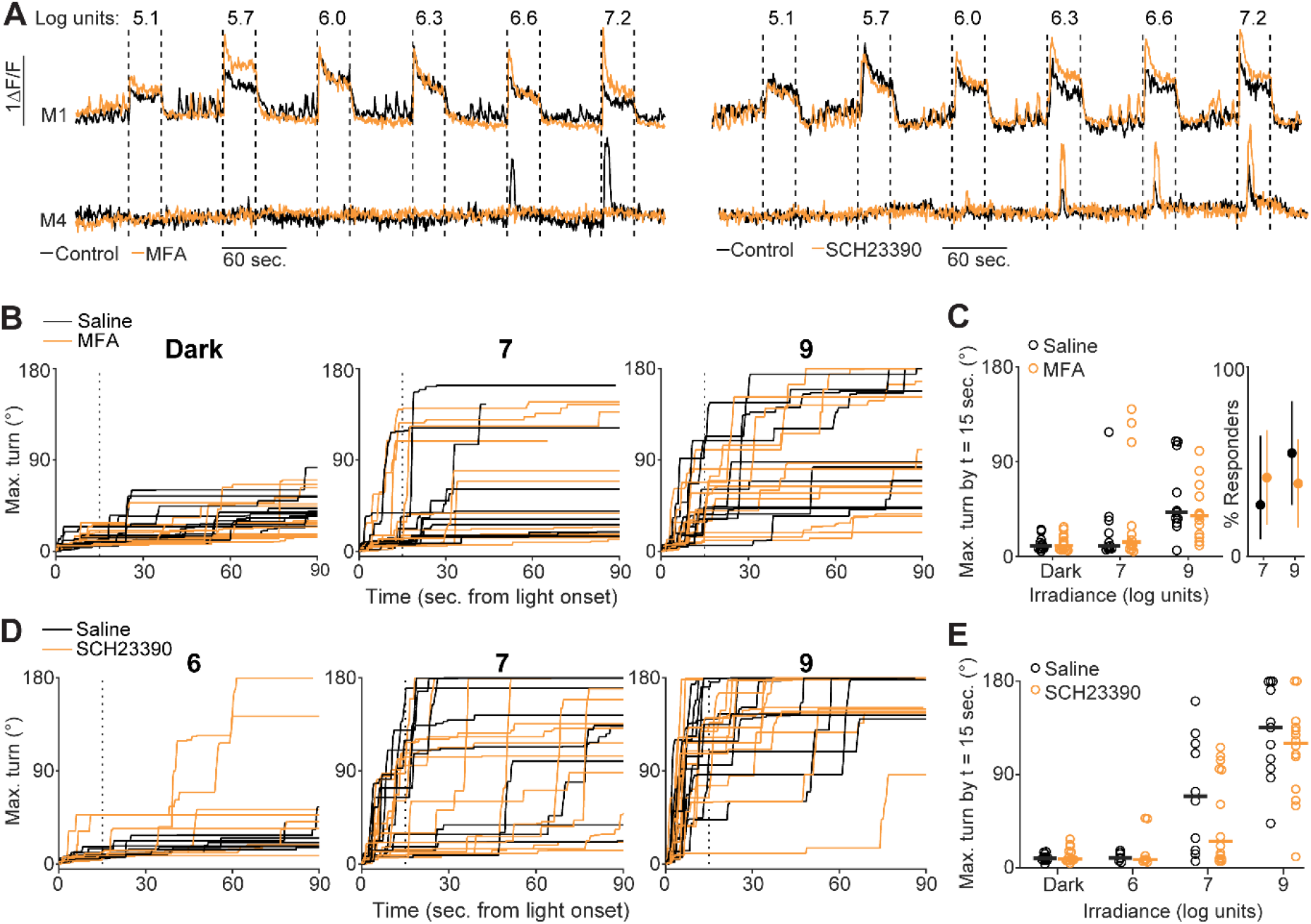
Photoaversion does not require gap junction coupling in the retina. A. Ca^2+^ responses of M1 and M4 ipRGCs in the *ex vivo* retina to a range of light stimulus intensities (e.g., 7.2 log units = 10^7.2^ photons μm^−2^ s^−1^) before (black traces) and after (orange traces) incubation in MFA (meclofenamic acid, a gap junction antagonist; left panel) or SCH23390 (a type-1 dopamine receptor antagonist that increases the extent of gap junction coupling; right panel). Data are replotted from (Caval-Holme et al., 2019). B. Timeseries of maximum head turns of P7 mice that received a bilateral intraocular injection of either saline (black traces) or MFA (orange traces). Panels correspond to trials in darkness (left), trials with a 7-log unit stimulus, or trials with a 9-log unit light stimulus. Vertical dashed line: time samples (15 sec. after light onset) used for quantification in C. Some trials are not displayed due to head-tracking errors. C. Comparison of head turns between the saline and MFA-injected mice in B. Left panel: maximum turn angle 15 seconds after stimulus onset. Horizontal bars indicate medians. Wilcoxon rank-sum test: p = 1.00, 0.73, and 0.60 for dark, 7 log unit, and 9 log unit trials, respectively. n = 11 saline-injected mice and 13 MFA-injected mice. Right panel: proportion of mice that responded to light with large head turns (right panel; a turn > 90° within 90 sec. of stimulus onset). Points: mean response rate; error bars: bootstrapped 95% confidence intervals. D. Timeseries of maximum head turns of P8 mice that received a bilateral intraocular injection of either saline (black traces) or SCH23390 (orange traces), as a function of irradiance. E. Comparison of head turns for the saline or SCH23390-injected mice in D. Horizontal bars indicate medians. Wilcoxon rank-sum test: p = 0.37, 1.00, 0.75, and 0.85 for comparisons of head turns on dark, and 6 log unit, 7 log unit, and 9 log unit trials, respectively. n = 11 saline-injected mice and 14 SCH23390-injected mice.

We also performed bilateral intraocular injections of the type-1 dopamine receptor antagonist SCH23390 (~14 μM in the vitreous humor). This manipulation increases light sensitivity, primarily in M2-M6 ipRGCs, via an increase in gap junction coupling (Caval-Holme et al., 2019). Mice injected with SCH23390 exhibited similar photoaversion to saline-injected littermate controls (**Figures 4D, 4E**). If M2-M6 contribute to photoaversion at lower light intensities, we hypothesized that mice injected with SCH23390 would respond to a 10^6^ photons μm^−2^ s^−1^ light stimulus (below the light intensity threshold for photoaversion in WT mice). However, mice who received SCH23390 did not display an avoidance response to this light intensity (**Figure 4E**).

### Connexin expression, role in ipRGC photosensitivity, and contribution to photoaversion

To complement our pharmacological approach, we used mice that lack specific connexin proteins to assess the impact of disrupted gap junction coupling amongst ipRGCs on photoaversion. In adults, ipRGCs express Cx30.2 (Meyer et al., 2016) and Cx36 (Harrison et al., 2021). To identify the connexins expressed in ipRGCs in development, we performed immunohistochemical analyses of retinas of mice from three transgenic lines in which regulatory elements of the Cx30.2 (Cx30.2^lacZ/+^) (Kreuzberg et al., 2006), Cx36 (Cx36^lacZ/+^) (Deans et al., 2001), and Cx45 (Cx45^fl/+^) (Maxeiner et al., 2005) genes drive expression of a reporter in place of the endogenous coding sequence (**Figure 5A**; see **Methods**). We crossed the Cx30.2 and Cx36 mutant mice with Opn4::eGFP mice and looked for co-localization of β-galactosidase and GFP. We found that roughly half of GFP+ ipRGCs in the retinas of Cx30.2^lacZ/+^; Opn4::eGFP mice expressed β-galactosidase (**Figure 5B, 5C**). In Cx36 ^lacZ/+^; Opn4::eGFP retinas, β-galactosidase expression was sparse in the ganglion cell layer as reported previously (Hansen et al., 2005) and rarely colocalized with GFP+ ipRGCs. Note, β-galactosidase was expressed in numerous cells within the inner nuclear layer, presumably in AII amacrine cells, indicating that this reaction product accurately reflects Cx36 expression at this age (data not shown).

**Figure 5:**
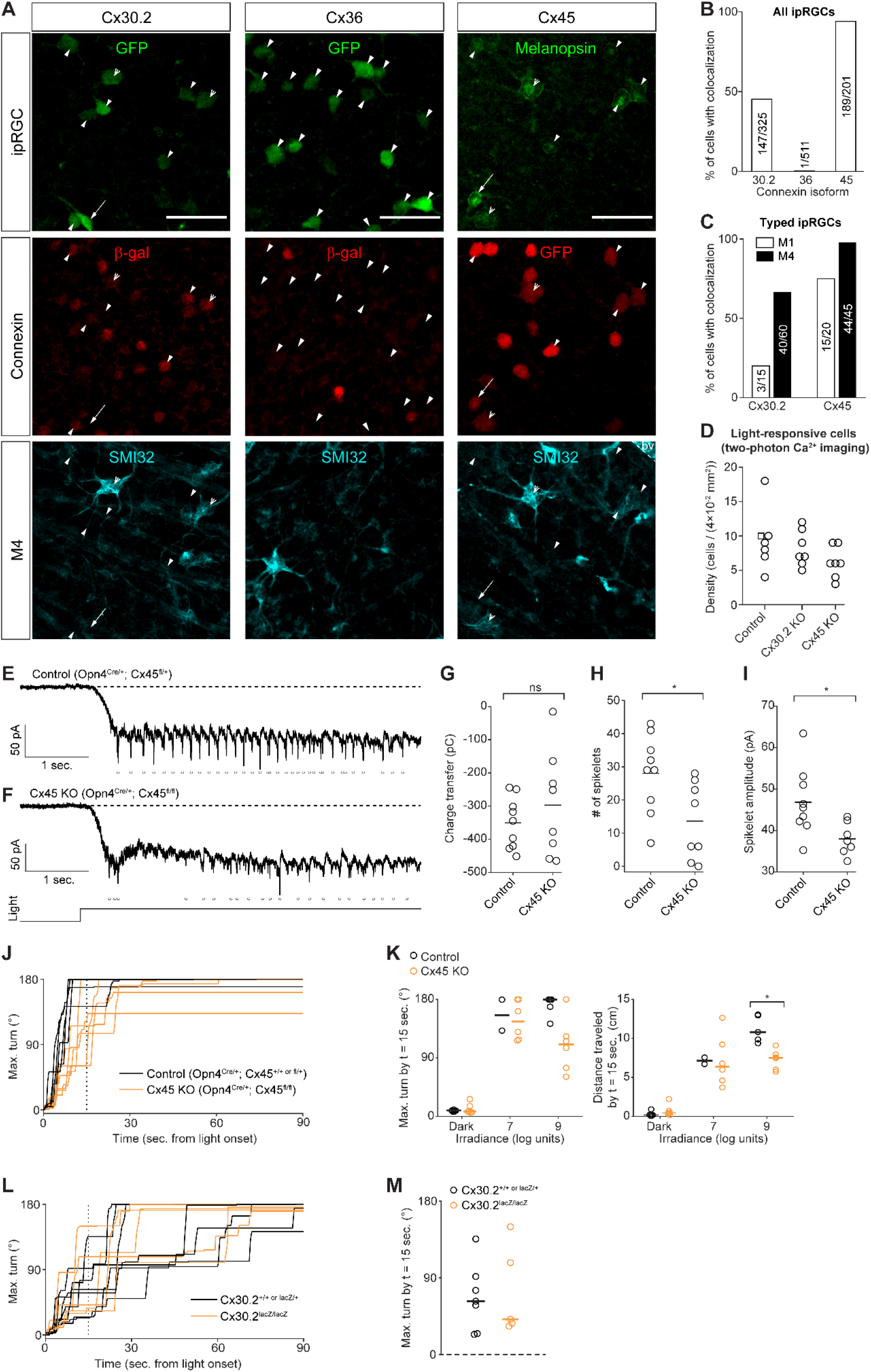
Connexin expression, role in ipRGC photosensitivity, and contribution to photoaversion. A. Immunohistochemistry images indicating expression of Cx30.2 (left column), Cx36 (middle column), and Cx45 (right column) in ipRGC types. Connexin expression was visualized using transgenic mouse lines (Opn4::eGFP Cx30.2^lacZ/+^, Opn4::eGFP Cx36^lacZ/+^, and Opn4^Cre/+^ Cx45^fl/+^) that express *lacZ* and *GFP* in place of the genes encoding Cx30.2, Cx36, and Cx45, respectively. Open arrowheads: M4 ipRGCs (GFP+ or melanopsin+ and SMI32+). Arrows: M1 ipRGCs (dendritic stratification in the outermost OFF layer of the inner plexiform layer). Closed arrowheads: ipRGCs of unknown type. Scalebar: 50 μm. bv: blood vessel labeled by the anti-mouse secondary antibody used to detect SMI32 (see Methods). B. Bar graph indicating the proportions ipRGCs with colocalized markers for connexin expression. The number of cells is indicated within each bar. N = 2 Opn4::eGFP Cx30.2^lacZ/+^ mice, 2 Opn4::eGFP Cx36^lacZ/+ or lacZ/lacZ^ mice, and 3 Opn4^Cre/+^ Cx45^fl/+^ mice. C. Same as B, but for ipRGCs identifiable as M1 or M4. D. The density of light-responsive cells in *ex vivo* retinas was not significantly affected by genetic deletion of Cx30.2 or Cx45. Each point represents one field of view (FOV) from a two-photon population Ca^2+^ imaging experiment. Light stimuli (λ_max_ = 420 nm) were a series of 30 sec. pulses from 3.7 to 7.2 log units, identical to those used in (Caval-Holme et al., 2019). Control: retinas from wild type mice of genetic backgrounds matched to the Cx30.2 KO and Cx45 KO. N = 6 fields of view (FOV) from 2 Opn4::eGFP mice and 1 Opn4^Cre/+^ Cx45^fl/+^ mouse (square symbol; note, one copy of the Opn4 gene is replaced with Cre, as in the Cx45 KO). Cx30.2 KO: 7 fields of view from 2 mice. Cx45 KO: 7 fields of view from 3 mice. One-way ANOVA (F = 1.88, 2 degrees of freedom, p = 0.18). E. Voltage-clamp recording of light-evoked current in an M4 ipRGC in the *ex vivo* retina of a Cx45 heterozygote (Opn4^Cre/+^ Cx45^fl/+^) mouse. Dashed line: holding current reference. Points: detected spikelets (inward currents magnitude > 30 pA). Illumination (λ_max_ = 420 nm, irradiance = 10^8^ photons μm^−2^ s^−1^) lasted for 5 sec. Light monitor below panel F. Methods for spikelet recording: Cells were held at −80 mV. The internal solution contained QX314 (5 mM), a voltage-gated sodium channels antagonist. Recording solution contained 50 μM D-AP5, 20 μM DNQX, and 8 μM DHβE. See (Caval-Holme et al., 2019) for further details. F. Same as E. except the cell was from a littermate Cx45 knockout (Opn4^Cre/+^ Cx45^fl/+^) mouse Recorded on the same day as the cell in E. G. Comparison of light-evoked charge transfer (pico-coulombs) in M4 ipRGCs from Cx45 Hets and Cx45 KOs. Horizontal lines: medians. ns: not significant, Wilcoxon rank-sum test p = 0.67. Cx45 Het: n = 9 cells from 4 mice. Cx45 KO: n = 8 cells from 3 mice. H. Comparison of the number of light-evoked spikelets in M4 ipRGCs from Cx45 Hets and Cx45 KOs. Horizontal lines: medians. *p < 0.05 Wilcoxon rank-sum test. Same cells as in G. I. Comparison of the amplitude of light-evoked spikelets in M4 ipRGCs from Cx45 Hets and Cx45 KOs. Horizontal lines: medians. *p < 0.05 Wilcoxon rank-sum test. Same cells as in G. and H., with one Cx45 KO cell excluded as it had no spikelets. J. Timeseries of maximum head turns of P8 Control (Opn4^Cre/+^ Cx45^+/+^ (blue traces) or Opn4^Cre/+^ Cx45^fl/+^ (red traces)) and Cx45 KO (red traces; Opn4^Cre/+^ Cx45^fl/fl^) mice. K. Comparison of head turns (left) and travel distance (right) in P8 genetic control (Opn4^Cre/+^ and Opn4^Cre/+^ Cx45^fl/+^ littermates) and Cx45 KO (Opn4^Cre/+^ Cx45^fl/fl^) mice. Control: n = 3 Opn4^Cre/+^ and 2 Opn4^Cre/+^ Cx45^fl/+^. Cx45 KO: n = 6. *p < 0.05, Wilcoxon rank-sum test. Horizontal bars indicate medians. L. Timeseries of maximum head turns of P7 Control (black traces; Cx30.2^+/+ or lacZ/+^) and Cx30.2 KO (red traces; Cx30.2^lacZ/lacZ^) mice. Light stimulus intensity was 10^9^ photons μm^−2^ s^−1^. M. Comparison of the maximum head turn 15 sec. after light onset. Horizontal bars indicate medians. Same mice as A. N = 7 Control mice and 5 Cx30.2 KO mice.

A third connexin, Cx45, has been observed in RGC types though not previously associated with ipRGCs (Blankenship et al., 2011; Hansen et al., 2005). To assess the expression of Cx45 in ipRGCs in the neonatal retina, we used an intersectional transgenic strategy in which we crossed Opn4^Cre^ mice with a mouse line in which Cx45 regulatory elements drive expression of eGFP in place of the Cx45 open reading frame (Maxeiner et al., 2005). Thus, in Opn4^Cre/+^ Cx45^fl/+^ retinas, ipRGCs will express both GFP and Cx45 (**Figures 5A**). Together, these data indicate neonatal ipRGCs express primarily Cx45, to a lesser extent, Cx30.2, and very little Cx36.

We next assessed the effect of Cx30.2 and Cx45 deletion on the light responses of ipRGCs. In Cx30.2^lacZ/lacZ^ (Cx30.2 KO) and Cx45 KO retinas, two-photon calcium imaging revealed light-responsive cells at densities that were not significantly different from genetic controls (**Figure 5D**), suggesting that gap junction circuits are at least partially retained in both KO transgenic lines. As M4 ipRGCs almost universally expressed Cx45 (**Figure 5C**), we assessed the impact of Cx45 on gap-junction mediated light responses of M4 ipRGCs. Voltage-clamp recordings of light-evoked currents in M4 ipRGCs from Opn4^Cre/+^ Cx45^fl/fl^ retinas (Cx45 KO; in which GFP is present but Cx45 is knocked out) revealed a significant reduction in the number and amplitude of spikelets (**Figure 5E-I**) but not their elimination (Arroyo et al., 2016; Caval-Holme et al., 2019). Unfortunately, we could not generate a Cx30.2/45 double KO with breeding because these two genes are located on the same chromosome with an estimated recombination frequency of < 4%.

Finally, we assessed photoaversion behavior in Cx30.2 KO and Cx45 KO mice (**Figure 5J-M**). At threshold levels of light, Cx30.2 KO and Cx45 KO mice had photoaversion behavior similar to their heterozygous littermates. Cx45 KO mice exhibited a small but statistically significant decrease in light-evoked movement at saturating light levels (**Figure 5J, 5K**), indicating that gap junction networks may contribute within this saturating stimulation regime. Because M1 ipRGCs exhibit minimal gap junction coupling in the neonatal retina (Caval-Holme et al., 2019), these results are consistent with M1 ipRGCs providing the principal drive for photoaversion.

### M1 ipRGCs are weakly depolarized by retinal waves

At the same age that there is photoaversion, the retina also exhibits retinal waves (**Figure 6A**) (Blankenship and Feller, 2010; Feller et al., 1996, 1997). These retinal waves are mediated by the diffuse release of excitatory neurotransmitters and are therefore thought to depolarize most cells in the ganglion cell layer, including all RGCs (Ford et al., 2012), including ipRGCs (Kirkby and Feller, 2013). Action potentials generated by retinal waves have been detected in the superior colliculus, lateral geniculate nucleus and visual cortex (Ackman et al., 2012; Mooney et al., 1996; Weliky and Katz, 1999). However, neonatal mice move little in the absence of light stimuli (Delwig et al., 2013; Johnson et al., 2010), indicating that the motor movements associated with photoaversion are selectively triggered by light-evoked neural activity. How does the neonatal mouse brain distinguish light-induced bursts of action potentials in ipRGCs from those induced by retinal waves?

**Figure 6:**
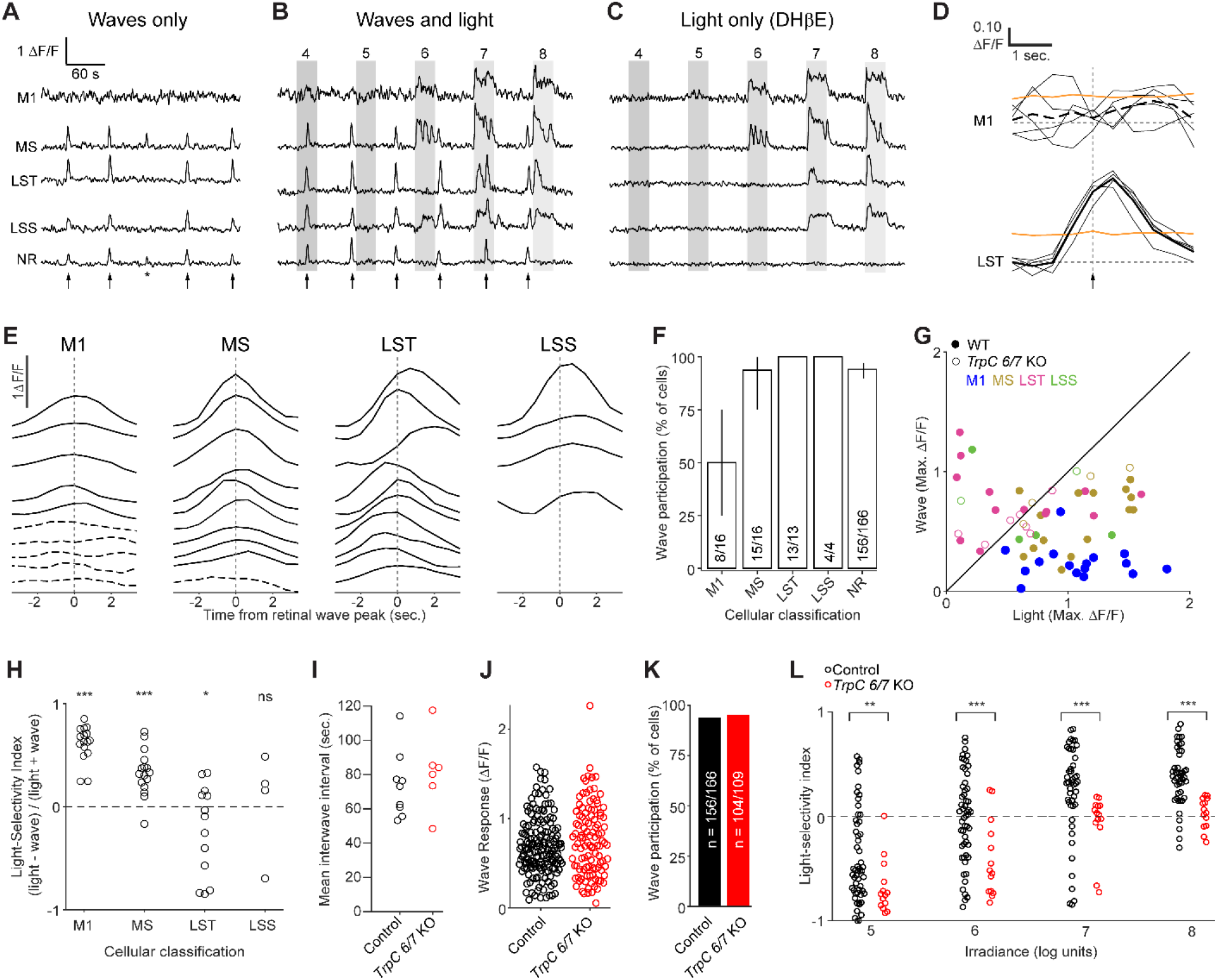
M1 ipRGCs are weakly depolarized by retinal waves. (A-C). Example ΔF/F traces from the same cells under a series of imaging conditions: A. waves only, B. waves and light stimulation, and C. light stimulation only (with waves blocked by DHβE). Arrows indicate timing of waves. Asterisk indicates a wave that involved only a portion of the field of view and was excluded from the analysis. Grey boxes indicate duration of light stimuli (λ_max_ = 470 nm; 10^4^-10^8^ photons μm^−2^ s^−1^). Waves-only and light-only traces were analyzed further. MS, LST, and LSS: functional cluster assignments based on each cell’s light response (**Figure 2**). NR: a randomly selected neuron within the ganglion cell layer that was not responsive to light. D. Sample wave-triggered averages for two of the cells in (A-C). Thin black lines: epochs used to generate the wave triggered average (thick black lines). Orange lines: 95^th^ percentiles of distributions of averages triggered from random non-wave locations. Vertical dashed line indicates the timing of the population wave peak. Top: M1 cell (average trace shown as dashed line) did not exceed the 95^th^ percentile at any point and was therefore defined as exempt from waves. Bottom: the LST cell clearly participates in waves. E. Representative wave-triggered average traces, vertically offset for clarity. Cells are a subset of those in Figure 2. Dashed lines indicate a lack of wave participation. M1: anatomically identified M1 ipRGCs. F. Proportions of light responsive (M1, MS, LST, and LSS) and non-responsive (NR) cells that participated in waves. N = 9 imaging fields of view from 4 mice (1:1 C129:C57BL6/J or Opn4::eGFP). Cell counts indicated within the bar chart. Error bars are bootstrapped 95% confidence intervals. G. Maximum ΔF/F responses of individual ipRGCs to waves plotted against maximum responses to light (10^7^ photons μm^−2^ s^−1^). Each point represents one cell and is color-coded according to functional cluster assignment, based on the cell’s light response (Figure 2). Ns for cells from Control mice (closed circles) are given in F. For *TrpC 6/7* KO (open circles), n = 1 M1, 4 MS, 8 LST, 2 LSS cells from 5 imaging fields of view from 2 mice. H. Light selectivity index (calculated as 1 - (W/L), L being the maximum ΔF/F response of a cell to light stimulus and W being the maximum response to waves) for cells from the functional clusters defined in Figure 2. Light intensity: 10^7^ photons μm^−2^ s^−1^. Only the light-responsive cells from wildtype mice are shown. Student’s t-test with Bonferroni correction. *p < 0.05, ***p < 0.0005, ns: not significant (p = 0.22). Ns are given in B. and D. I. Average inter-wave time intervals from different fields of view in retinas from Control and *TrpC 6/7* KO mice. Two sample t-test p = 0.52. J. The distributions of ΔF/F responses to retinal waves were not significantly different between non-light-responsive cells from Control and *TrpC6/7* KO retinas (two sample t-test p = 0.13). K. The proportion of cells participating in retinal waves was indistinguishable between non-light-responsive cells from Control and *TrpC6/7* KO retinas (χ^2^ test, p = 0.61). L. Light selectivity index in Control (1:1 C129:C57BL6/J; n = 9 fields of view from 3 mice) and *TrpC 6/7* KO (n = 6 fields of view from 2 mice) retinas, at four different light intensities. Wilcoxon rank-sum test with Bonferroni correction: **p < 0.005, ***p < 0.0005.

To test whether the ipRGCs driving photoaversion are strongly activated by retinal waves, we used two-photon Ca^2+^ imaging. We compared the Ca^2+^ transients under three conditions: 1) waves without light responses (**Figure 6A**); 2) waves with light responses (**Figure 6B**); 3) light responses without waves (**Figure 6C**). Each cell’s participation in retinal waves was computed from a retinal wave-triggered average (**Figure 6D, E**). Cells whose retinal wave-triggered averages were significantly larger than noise fluctuations (determined by bootstrapping non-wave fluorescence changes; see Methods) were classified as participating in retinal waves. Finally, we functionally classified cells based on their light-evoked Ca^2+^ transients, and independently identified M1s from their anatomy, as described above. While all LST cells (20/20) and LSS cells (5/5) and most MS cells (15/16) participated in retinal waves, we found that only 50% of anatomically identified M1s (8/16 in total, with 2/6 identified via anti melanopsin immunohistochemistry and 6/10 identified in *Opn4::eGFP* retinas) participated (**Figure 6F**; For a representative imaging session, see http://fellerlab.squarespace.com/movies). M1s thus had strikingly low rates of wave participation relative to other light-sensitive cells (> 98% of MS, LST, and LSS) and the broader population of light-insensitive cells (~94%; 156/166 cells). Together, these results indicate that M1 ipRGCs are among the few cells in the GCL that are weakly depolarized by retinal waves.

We next compared the size of depolarizations induced by waves to those induced by light stimulation by comparing the magnitude of Ca^2+^ responses (ΔF/F; **Figure 6G**). To test for differences in wave and light-evoked responses in cells of each functional cluster, we defined a light selectivity index (the ratio of the Ca^2+^ response evoked by light less the Ca^2+^ response evoked by waves, divided by their sum). Indeed, M1/HS and MS cells had smaller response to waves than light, in contrast to LST, which had the opposite, and LSS cells which had similar responses (**Figure 6H**). While the properties of retinal waves in the *TrpC 6/7* KO were indistinguishable from control (**Figure 6I-K**), light-responsive cells had significantly smaller light selectivity indices at all light intensities (**Figure 6L**), consistent with the loss of photosensitivity we observed in M1/HS cells (**Figure 2**). Thus, M1 ipRGCs exhibit larger Ca^2+^ responses to light than retinal waves and this functional specialization is disrupted in the *TrpC 6/7* KO.

## DISCUSSION

Photoaversion is an innate behavior that is present early in development, before maturation of the circuits that mediate image-forming vision. Here we report that the behavior is reduced in mice lacking the TRPC6/7 phototransduction mechanism and is normal in mice with reduced gap junction coupling and in mice in which all ipRGCs other than the Brn3b-negative M1s are ablated. Two-photon calcium imaging in isolated retina confirm that TRPC6/7 KO reduces light responses specifically in M1 ipRGCs. In addition, we discovered that M1s are weakly depolarized by retinal waves, providing a potential cue by which the brain distinguishes M1 depolarizations induced by light from those induced by retinal waves. Together these data indicate that M1s provide a dedicated pathway for photoaversion during development.

### Brn3b-negative M1 ipRGCs are the principal drivers of photoaversion

Our data are consistent with a functional organization in which M1 ipRGC, specifically the subset lacking expression of the Brn3b transcription factor, drive photoaversion. The evidence for this is several-fold: First, we found that the threshold light intensity required to elicit photoaversion is bright enough to activate M1 and M3 ipRGCs (note: M3s do not tile the retina with their dendritic arbors, suggesting they may not constitute a defined RGC type (Berson et al., 2010; Schmidt and Kofuji, 2011)) but too dim to activate M2, M4, M5, and M6 ipRGCs (note: the light intensity threshold for M4s is low in the adult (Sonoda et al., 2018)). Second, genetic deletion of TRPC6 and TRPC7 ion channels, which mediate light responses primarily in M1 ipRGCs in the adult (Jiang et al., 2018; Sonoda et al., 2018), dramatically reduced photoaversion and diminished photosensitivity in *ex vivo* M1s but had little effect on photosensitivity in other ipRGCs. Third, Opn4^Cre^ Brn3b^DTA^ mice in which all ipRGCs save for ~400 Brn3b-M1s are ablated, had apparently normal photoaversion. Fourth, intraocular injections of drugs that modulate the extent of gap junction coupling, primarily affecting the light sensitivity of M2-M6 ipRGCs (Caval-Holme et al., 2019), had no effect on photoaversion. Similarly, genetic knockout of connexins expressed in ipRGCs did not affect photoaversion at threshold light intensities. Finally, we found that M1s have limited depolarizations by retinal waves, a property that may make them uniquely well-suited to the requirements of photoaversion behavior.

Do M2-M6 contribute to photoaversion at saturating intensities? While *TrpC 6/7* KO mice exhibited no response to a near-threshold light intensity they made small head turns in response to saturating intensities (**Figure 1**). This observation implies either a contribution from M2-M6 or residual signaling from M1s, consistent with small Ca^2+^ (**Figure 2**) and current (Jiang et al., 2018) responses measured in *TrpC 6/7* KO M1s. Mice lacking Cx45 in ipRGCs (**Figure 5**) exhibited reduced spikelets in M4 ipRGCs and slightly reduced photoaversion at saturating intensities (Note: that in mice that had received an intraocular injection of MFA there was no detectable reduction in photoaversion. This is likely due to highly variable behavior in mice that receive intraocular injections, which leads to irregular and prolonged pupillary constriction).

### Identification of connexins in developing ipRGCs

In the adult retina, the coupling patterns of ipRGCs are well-described. IpRGCs are coupled to several types of amacrine cells (Müller et al., 2010; Pottackal et al., 2021; Reifler et al., 2015), many of which fire action potentials (Reifler et al., 2015). Coupling between ipRGCs has not been observed. Cx36 has been implicated in ipRGC-AC coupling (Harrison et al., 2021) and a subset of ipRGCs express Cx30.2 (Meyer et al., 2016). Cx45 is only thought to be present in bistratified RGCs in the adult (Schubert et al., 2005), and thus potentially could be present M3s.

The coupling pattern and connexin expression during development is distinct from the adult. M2-M6 ipRGCs are extensively coupled during development and this strongly contributes to the light response (Arroyo et al., 2016; Caval-Holme et al., 2019). In contrast to the adult (Harrison et al., 2021), we found little evidence for Cx36 expression in ipRGCs during development (**Figure 5**), consistent with previous work from our lab showing Cx36 expression in the ganglion cell layer is low during the first postnatal week (Hansen et al., 2005). We found a small percentage of ipRGCs to express Cx30.2 (**Figure 5**), consistent with coupling described in the adult. Finally, we discovered that ipRGCs express Cx45 during development. It is important to note that Cx45 expression was determined using an intersectional transgenic approach – thus expression of Cx45 at any point in development will lead to expression of GFP. In contrast, Cx30.2 and Cx36 expression was based on expression of β-galactosidase which will reflect the level of expression only at the age at which we did the experiment. That said, Cx45 KO mice had a physiological phenotype, with M4-ipRGCs displaying fewer spikelets than observed in control animals (**Figure 5E, 5F**). This provides a potential basis for M4s contributing to photoaversion at saturating intensities.

### Why don’t retinal waves interfere with photoaversion?

Previously, it was assumed that all RGCs, including ipRGCs, participate in waves due to their ubiquitous expression of nicotinic acetylcholine receptors (Aizenman et al., 1990; Zoli et al., 1995) and the volume release of acetylcholine by burst amacrine cells (Ford et. al. 2012). However, a recently published single-cell transcriptomics dataset revealed that M1 ipRGCs express strikingly low levels of genes encoding nicotinic acetylcholine receptor subunits (Tran et al., 2019). We found that M1 ipRGCs exhibited depolarizations induced by light stimulation that were stronger than those induced by retinal waves, with about half of anatomically identified M1 ipRGCs exhibiting no detectable calcium transient in response to waves (**Figure 6**).

In addition to the strength of depolarization, there are other differences between retinal wave-evoked and light-evoked activity in RGCs that could be utilized by the ipRGC-recipient regions of the brain to discriminate the two sources of input. Light stimuli induce synchronous depolarizations lasting for 10s of seconds while retinal waves induce propagating depolarizations lasting 1-2 seconds per RGC (Ford et al., 2012; Meister et al., 1991). Light stimuli can also activate ipRGCs across the entire retina, while retinal waves have finite propagation (Feller et al., 1996). Though we have not explored the full temporal and spatial properties of visual stimuli that evoke photoaversion, M1s integrate photons over large ranges of stimulus duration and generate prolonged responses to brief light stimuli (Emanuel et al., 2015), including single photon absorptions (Do et al., 2009). Hence duration of depolarization and the number of simultaneously depolarized neurons may be other features that allow the brain to differentiate light-evoked and spontaneous activity. Nevertheless, our observation of strikingly weak responses of M1 ipRGCs to waves suggests that the nervous system has evolved to minimize inputs from spontaneous activity to the information channels subserving photoaversion. Exemption from retinal waves may also underlie the absence of retinotopic and eye-specific refinement observed in retinorecipient regions receiving predominantly M1 projections (Ecker et al., 2010), increasing the extent of light’s spatial integration in non-image-forming pathways.

In summary, we conclude that M1 ipRGCs constitute a distinct information channel that mediates photoaversion during the period of early postnatal development when retinal waves drive activity-dependent refinement of retinal projections to the brain.

## AKNOWLEDGEMENTS

F.C.-H. was supported by F31EY028022-03. M.B.F. was supported by NIH RO1EY019498, NIH RO1EY013528, and NIH P30EY003176. M.L.A. and T.M.S. were supported by 1DP2EY027983. L.B. was supported by the NIH Intramural Research Program (project ZO1-ES-101684). Confocal images were obtained on a microscope at the Molecular Imaging Center at UC Berkeley. Finally, we thank Michael Tri Hoang Do and members of the Feller Lab for their comments on the manuscript.

## AUTHOR CONTRIBUTIONS

F.C.-H., T.M.S., and M.B.F. designed experiments and wrote the manuscript. F.C.-H., M.L.A., A.T., A.Q.C., and Y.Z. performed experiments. A.Q.C. analyzed participation of ipRGCs in retinal waves. F.C.-H., A.Q.C., and Y.Z. analyzed photoaversion data. B.S. and F.C.-H. created mouse tracking algorithms and built the behavior apparatus. L.B. originally generated *TrpC* KO mice.

## METHODS

### RESOURCE AVAILABILITY

#### Lead contact

Further information and requests for resources and reagents should be directed to and will be fulfilled by the Lead Contact, Marla Feller.

#### Materials availability

This study did not generate new unique reagents.

#### Data and code availability

- The photoaversion behavior and two-photon calcium imaging datasets have been deposited, in a preprocessed format, at Mendeley and are publicly available as of the date of publication. DOIs are listed in the key resources table. Electrophysiology and immunohistochemistry datasets are available upon request.
- All original code has been deposited at https://github.com/FellerLabCodeShare and https://github.com/Llamero/Mouse_tracking_macro and is publicly available as of the date of publication. DOIs are listed in the key resources table.
- Any additional information required to reanalyze the data reported in this paper is available from the lead contact upon request.

### EXPERIMENTAL MODEL AND SUBJECT DETAILS

Animal procedures were approved by the University of California, Berkeley Institutional Animal Care and Use Committees and conformed to the National Institutes of Health Guide for the Care and Use of Laboratory Animals, the Public Health Service Policy, and the Society for Neuroscience Policy on the Use of Animals in Neuroscience Research. Photoaversion assays were performed on P7 and P8 mice of either sex. Animal health was monitored daily, and only healthy animals were used in experiments. WT mice were from the C57BL/6J strain. Melanopsin knockout mice were generated by crossing Opn4^Cre/+^ mice, in which one of the copies of melanopsin is replaced by Cre to generate Opn4^Cre/Cre^ homozygotes. *TrpC 6/7* knockout mice were generated from *TrpC 3/6/7* knockout mice obtained from Professor Tiffany Schmidt (Northwestern University). As *TrpC* knockout mice were generated on a 1:1 C57BL/6J:C129 genetic background, 1:1 C57BL/6J:C129 wild-type mice served as controls for experiments with *TrpC* 6/7 knockouts.

### METHOD DETAILS

#### Photoaversion assay apparatus

The behavior chamber consisted of an open-topped transparent plastic box (Warner Instruments) measuring 10.8 × 3.5 × 2.5 cm (L × W × H). The chamber was painted on all surfaces (except the inner and outer faces at one end, left transparent for light entry) with matte black low-aerosol spray paint. Matte black construction paper was cut into a rectangle and placed on the chamber floor to provide traction and minimize optical reflections that interfered with automated tracking of mice, with a fresh piece of paper used for each behavioral session. The base of the chamber was warmed to 35± 2° Celsius by a heating pad, and the temperature was monitored continuously with a thermistor probe (Warner Instruments).

Video monitoring of the behavior was achieved with a camera (Thorlabs CS165MU with Thorcam software) fitted with a high-pass optical filter (Thorlabs FEL0600) to prevent light from the stimulus LED from reaching the detector. The camera, along with an infrared LED light source (Thorlabs M970L4, λ_max_ = 970 nm) fitted with a spherical lens, was mounted on a pole and focused on the chamber from 45 cm above the floor of the chamber. Video frames at a resolution of 32 × 105 pixels were collected at 27.5 frames/second.

The light stimulus was delivered by an LED (Luxeon SP-05-B4 LED module; λ_max_ = 470 nm; full width at half-maximum = 15 nm) coupled to collimating optics (Thorlabs). The timing and intensity of light stimulation were controlled with pulse width modulation, using an Arduino Uno running custom software written in the Arduino programming language. The onset of a light stimulus also triggered a brief flash from an infrared LED (LED Supply L3-0-IR5TH50-1, λ_max_ = 850 nm) mounted within the field of view of the camera but out of the line of sight of the mouse, thus recording the timing of the light stimulus within the video of each behavioral trial (the infrared light also flashed at the t of the dark trials). Irradiance measurements were obtained by converting measurements of optical power to photon flux. Measurements of optical power were constant along the length of the chamber, as expected for a minimally diverging light beam.

#### Photoaversion assay

Mice were dark-adapted in the home cage for one hour before experiments. During a behavioral session, a mouse pup was removed from the home cage, and an infrared-reflective sticker was affixed to its head. The center of the sticker was centered between the ears with respect to the rostral-caudal body axis. The mouse pup was then acclimated to the chamber for four minutes. Before each trial, the mouse was placed in the chamber with its head angled to within +=30° of the stimulus source (the axis of the collimated light beam from the LED; Note: we collimated the LED light after observing in pilot experiments using point-source stimuli (the LED lens only) that the magnitude and dynamics of the turning response varied with the distance of the mouse to the source, as expected for a highly-diverging beam).

Video recording commenced and the pup was monitored for 30 seconds in the dark. If the pup turned so that its head angle exceeded ± 45° from the LED, it was repositioned, and the trial reted. A maximum of three rets were allowed before a pup would be excluded from the study. When individual pups were tested on repeated trials that included light stimuli, they were left for 60 seconds in the dark between trials to allow the ipRGCs to partially recover from light adaptation (Do and Yau, 2013). In pilot experiments, we observed that the magnitude and dynamics of the light-evoked turning behavior were similar for up to three trials at the saturating irradiance (9 log units; data not shown). After a behavioral session, the chamber was cleaned with 70% ethanol, which was allowed to evaporate for four minutes before commencing a session with the next mouse. To minimize differences in the behavior due to circadian variability, we performed experiments only during the subjective day portion of the 12/12 hour light/dark cycle in the rodent housing rooms.

For behavioral experiments measuring the intensity-dependence of photoaversion (**Figure 1**), each mouse pup was tested at a single light stimulus intensity. In some mice, a trial in the dark was recorded preceding the trial with a light stimulus. For behavior sessions involving transgenic mice and eye injections, a session for one mouse consisted of 90-second trials, ting with a trial in the dark, and then proceeding with trials with light stimuli in order of increasing irradiance.

#### Open field test

To measure movement without activating photoaversion pathways, we performed an open field test in which we placed a P11 pup into a room temperature (20°C) Pyrex beaker with an open top. Lighting conditions were dim enough not to evoke a photoaversion response. Behavioral recordings were made using the same camera as in the *Photoaversion behavior apparatus* section, with a resolution of 381 × 365 pixels at 15 frames/second.

#### Enucleation

Enucleation procedures were conducted just after birth (P0). A maximum of 1 hour of nursing was allowed before surgery. Pups were anesthetized with an IP injection of a mixture of ketamine (40 mg/kg) and xylazine (5 mg/kg), then placed on ice for 1 to 4 minutes. A toe-pinch was performed to confirm the appropriate level of surgical anesthesia, after which the eyelid was cleaned using aseptic technique.

An incision was first made in each eyelid with a scalpel. The eye was lifted away from the orbit with forceps and severed from the optic nerve with surgical scissors. The eyelid was sealed with 0.5 μL to 1 μL of tissue adhesive (Surgi-Lock instant liquid tissue adhesive, Fisher Scientific, Pittsburgh, PA, USA). The procedure was then replicated on the other eye. Sham surgeries omitted the enucleation step but were otherwise identical.

After enucleation, the pups were immersed in lukewarm water bath for 30 seconds followed by an application of Lidocaine hydrochloride jelly USP, 2% (Akorn, Lake Forest, IL, USA and erythromycin ophthalmic ointment USP, 0.5% (Bausch & Lomb, Rochester, NY, USA) to prevent pain, swelling and infection. At the end of the procedure, Buprenorphine was administered and repeated 6-8 hours later. Pups were then evaluated the following day for additional need. In addition, NSAID (meloxicam) was given at the end of the procedure and repeated the following day. Pups were housed with their mother in a cage with nesting material before and after surgery. At P8, the pups were tested in the photoaversion assay.

#### Intraocular injections

Intraocular injections were performed as described previously. Drugs were dissolved in PBS and injected at concentrations of 500 μM (MFA) and 100 μM (SCH23390). Injections delivered 0.5 μL of solution into the vitreous cavity, which, at P7-8 contains a volume of ~3.5 uL (Kaplan et al., 2010). The concentrations of the drugs in the vitreous cavity would thus be reduced by a factor of ~0.14, to ~71 μM MFA and 14 μM SCH. Mice were recovered and dark adapted for ≥ 1 hour after injections.

#### Ex-vivo retina preparation

Mice were deeply anesthetized with isoflurane inhalation and euthanized by decapitation. Eyes were immediately enucleated, and retinas were dissected in oxygenated (95% O_2_, 5% CO_2_) ACSF (in mM, 119 NaCl, 2.5 KCl, 1.3 MgCl_2_, 1 K_2_HPO_4_, 26.2 NaHCO_3_, 11 D-glucose, and 2.5 CaCl_2_) at room-temperature under white light. In some experiments, 0.1 μM sulforhodamine 101 (SR101 Invitrogen) was added for visualization of vasculature. Each isolated retina was cut into two pieces. Each piece of retina was mounted over a 1–2 mm^2^ hole in nitrocellulose filter paper (Millipore) with the photoreceptor layer side down, dark-adapted for one hour, and transferred to the recording chamber of a two-photon microscope for imaging or electrophysiological recording. The whole-mount retinas were continuously perfused (3 ml/min) with oxygenated ACSF warmed to 32-34 °C by a regulated inline heater (TC-344B, Warner Instruments) for the duration of the experiment. Additional retina pieces were kept in the dark at room temperature in ACSF bubbled with 95% O_2_, 5% CO_2_ until use (maximum 8 hrs.).

#### Ex-vivo visual stimulation

Visual stimuli were delivered by LEDs (Thorlabs M420L2 or M470L2 λ_max_ = 420 or 470 nm; full width at half-maximum = 15 nm) coupled to a digital micromirror device (Digital Light Innovations Cel 5500). Stimuli were regularly calibrated with a radiometer (Newport). The intensity of the stimulus ranged from 10^4^ to 10^8^ photons μm^−2^ s^−1^, replicating the range of light intensities present at the retina from twilight to full sunlight (Allen et al., 2014). Each light stimulus lasted 30 sec. and was separated from the previous light stimulus by 60 sec. The stimuli had a 100% positive contrast (bright on dark background).

To decrease the background signal during the stimulus (due to emission by the phosphorescent objective glass and Cal 590, following absorption of the stimulus light), the stimulus was interleaved with imaging. the stimulus was delivered on the flyback of the fast axis scanning mirror during a unidirectional scan.

#### Two-photon calcium imaging

Each retina was cut into halves. To maintain control over light history, only one field of view was sampled from each half. Retina halves were bulk-loaded with the calcium indicator Cal 590 AM (AAT Bioquest) using a multicell bolus loading technique described previously (Blankenship et al., 2009; Stosiek et al., 2003). Two-photon fluorescence measurements were obtained with a modified movable objective microscope (MOM, Sutter instruments) using an Olympus 60X, 1.00 NA, LUMPlanFLN objective (Olympus America) for single cell resolution imaging (field of view: 203 × 203 μm). This MOM was equipped with through-the-objective light stimulation and two detection channels for fluorescence imaging. Two-photon excitation was evoked with an ultrafast pulsed laser (Chameleon Ultra II; Coherent) tuned to 1040 nm to image Cal 590 AM. Di-hydro-β-erythroidine (DHβE, 8 μM, Tocris) was added to the perfusion system immediately before imaging to block spontaneous retinal waves that would otherwise interfere with measurements of light responses. Previously, we showed that extended (≥ 1 hr.) blockade of waves with DHβE leads to an increase in the number of light sensitive cells (Arroyo et al., 2016). We carried out all experiments within the first 20 minutes of wave blockade, during which time the number of light-responsive cells does not increase (Caval-Holme et al., 2019). Laser power was set to 6.5 μW for imaging Cal 590 AM. The microscope system was controlled by ScanImage software (www.scanimage.org). Scan parameters were [pixels/line x lines/frame (frame rate in Hz)]: [256 × 256 (1.48 Hz)], at 2 ms/line.

#### Immunoassays

Whole-mount retinas were removed from the recording chamber and transferred to a 4% paraformaldehyde solution for 20 minutes at room temperature. Following fixation, retinas were washed in blocking buffer (1.5% BSA, 0.2% Na-Azide, 0.2% Triton X-100) (3 times, 10 minutes each time). Retinas were then incubated in a primary immunoreaction solution for 1-3 days at 4°C. Primary immunoreaction solution consisted of blocking buffer that contained 1:1000 rabbit anti-melanopsin (ATS Bio AB N38) and 1:250 mouse anti-SMI32 (Biolegend 801701). After incubation in the primary immunoreaction solution, retinas were washed in PBS (3 times, 10 minutes each time), and then incubated for 2 hours at room temperature in secondary immunoreactive solution containing one or more of the following secondary antibodies: 1:1000 donkey anti-rabbit conjugated to Alexa 488 (Invitrogen A21206) and 1:1000 goat anti-mouse conjugated to Alexa 647 (Invitrogen A21235). Retinas were washed again in PBS and then mounted on slides with an anti-fade agent (Vectashield H-1400, Vector Laboratories).

### QUANTIFICATION AND STATISTICAL ANALYSIS

#### Tracking of mouse head angle and position

After photoaversion assays, we tracked mouse head angle and position via automated tracking of a circular infrared-reflective sticker affixed to pups’ heads, using a custom algorithm implemented as a plugin in ImageJ (Note, neonates also produce ultrasonic vocalizations in response to bright light (Delwig et al., 2012). However, we found this to be more variable than movement). The fiducial was fabricated from a roll of infrared-reflective tape using a hole punch. A 16-gauge needle was used to punch a fiduciary hole near the edge of each sticker. Specifically, the circular fiducial was tracked in each movie frame using a Hough circle transform (HCT). While the HCT is very robust as segmenting circular objects in an image, it also adds an extra dimension to the search space corresponding to the circle radius and therefore is computationally intensive. We developed a modified HCT to track the circular fiducial with less computational time. After the circular fiducial is found in the first frame of the movie, the transform and search space in the following frames are narrowed down to a fiducial of similar radius and position. The algorithm was additionally optimized by multi-processing both the transform and search algorithms. More details on the HCT plugin can be found here: https://imagej.net/Hough_Circle_Transform.

To measure head orientation, a black dot was added to the inside edge of the circular fiducial. This allowed for extraction of orientation as a vector from the center of the black dot to the center of the circular fiducial. An ImageJ macro was written that used the HCT plugin to find the circular fiducial and segmented the black dot with the circle to obtain the vector orientation. The ImageJ macro used can be found here: https://github.com/Llamero/Mouse_tracking_macro. To quantify pups’ movements in the open field test, we tracked the base of the tail using deepLabCut (Nath et al., 2019).

#### Behavioral quantification

We wrote custom algorithms in MATLAB to preprocess and analyze the raw head angle and position data from the HCT tracking program and deepLabCut. Mouse head angle and position were filtered, to interpolate over brief (< 1 sec.) tracking errors. Turn angles were defined as the angular difference between the absolute head angle at time-sample t and the mean head angle within 0.5 sec. of the stimulus. We computed cumulative maximum from the turn angle dataset using MATLAB’s cummax function.

#### Image analysis of population calcium imaging movies

Movies were spatially median-filtered to remove high-frequency outlier noise and then registered relative to a frame in the middle of the movie to remove mechanical drift. The baseline movie frame (F_0_) was computed by taking the temporal median projection of all the movie frames. Each movie frame F. was normalized by dividing its difference from the baseline frame (F-F_0_) by the baseline frame ((F-F_0_)/F_0_) to produce a ΔF/F_0_ movie. Circular regions of interest (ROIs) were drawn on cells that displayed >20% increases in ΔF/F_0_ during at least one of the light stimuli. The ROIs and the ΔF/F_0_ movie was then imported into MATLAB for further analysis using custom algorithms. Traces for each ROI were computed as the mean value of the pixels enclosed by the ROI in each frame of the ΔF/F_0_ movie. For each ROI, the amplitude of the response to each light stimulus was computed as the difference between the peak ΔF/F_0_ value during the light stimulus and the ΔF/F_0_ value during the movie frame before the light stimulus. ROIs were defined as ‘light-responsive if the maximum amplitude of their ΔF/F_0_ signal during any light stimulus exceeded the mean ΔF/F_0_ in the 30 second interval preceding the light stimulus by more than six standard deviations.

#### Unsupervised clustering of light-evoked fluorescence transients

Fluorescence traces for all light-responsive cells were combined into a single matrix (cells x movie frames). The traces were high pass filtered at 0.01 Hz to remove slow changes in fluorescence caused by mechanical drift in the z axis and the samples corresponding to the 30 seconds preceding each light stimulus were removed. Each trace was normalized to its own maximum value. Functional clustering of light responsive cells from genetic background control mice was performed as described previously (Baden et al., 2016; Caval-Holme et al., 2019) and culminated in a set of sparse principal components weights and a gaussian mixture model. To classify light-responsive cells from TRPC 6/7 KO mice, we used the sPCA weights computed from the genetic background dataset to project their fluorescence traces onto the principal components. We then used the GMM from the genetic background dataset to assign the TRPC 6/7 KO cells to functional clusters.

#### Identification of ipRGC types

IpRGC types were identified at the conclusion of two-photon calcium imaging sessions using one of two strategies. In Opn4::eGFP mice, z stacks of GFP+ cells in the live retina were collected. In other mouse lines, ipRGCs were identified using melanopsin immunostaining. In the latter approach, fields of view in the fixed tissue were realigned to fields of view from two-photon calcium imaging by automated registration of common landmarks in the vasculature. Vasculature was visualized in live tissue using SR101 and in fixed tissue using an antibody against the neurofilament protein SMI32 that fortuitously labels blood vessels (see ***Immunoassays***). In both approaches, M1 ipRGCs were identified by their dendritic stratification at the outermost boundary of the in the inner plexiform layer, close to the inner nuclear layer (Caval-Holme et al., 2019).

#### Analysis of retinal waves

Inter-wave interval Population wave frames were identified by finding peak ΔF/F transients in the average trace of the whole field of view, using the MATLAB findpeaks functions with a minimum peak prominence of 1.5 after z-scoring. The difference between consecutive population wave locations was taken to be the inter-wave interval. Wave-triggered average ΔF/F traces, each spanning 11 frames, centered around population wave locations of the whole field of view, were averaged for each neuron to produce the wave-triggered average. A bootstrapping approach was used to generate a null distribution of randomly triggered averages. For each neuron with a given number of wave locations, the same number of virtual population wave locations were generated, such that the 11-frame wide window around the virtual wave location would not overlap with any of the actual wave windows. Averages were computed with the virtual locations in an otherwise identical manner and were computed with a new set of random wave locations 1000 times per neuron. A 95^th^ percentile array was computed with the MATLAB prctile function applied to this array of 1000 averaged traces. A neuron with a wave triggered average that at any point exceeded this 95^th^ percentile array was defined as participating in retinal waves and was otherwise defined as exempt from the retinal wave.

#### Statistical analyses

p < 0.05: *, p < 0.005: **, p < 0.0005: ***. Data that significantly violated the assumptions of parametric statistical tests were assessed using non-parametric tests. Wilcoxon-rank sum tests were used to compare medians of independent samples. Error bars represent bootstrapped 95% confidence intervals. Horizontal bars indicate medians. The number of replicates and statistical results are provided in figure captions.

